# Penton-dodecahedron of fowl adenovirus serotype 4 as a vaccine candidate for the control of related diseases

**DOI:** 10.1101/394072

**Authors:** Qiuxia Tang, Ruyi Dangi, Li Qiu, Zengqi Yang, Xinglong Wang

## Abstract

In some serotypes of adenovirus (Ad), the penton base protein and attached trimeric fiber assemble into dodecameric virus-like particles called penton-dodecahedron (Pt-Dd), which can internalize into cells and can be used to deliver the vaccine antigen and drugs. Fowl adenovirus serotype 4 (FAdV-4) is an important poultry pathogens and causing seriously economic lost to poultry industry in China and several other counties. The produce of Pt-Dd in FAdV-4 infected cells as well as in those infected with the recombinant human Ad expressing fiber-1, fiber-2, and penton base was determine by Transmission electron microscopy (TEM). For the First time, we proved that FAdV-4 produced Pt-Dd in infected cells, which can also be assembled by the overexpressed recombinant proteins fiber-1, fiber-2, and penton base. Pt-Dd, as well as the recombinant proteins fiber-1, fiber-2, and penton base, were then used to immunize chickens. The humoral immune response, cell mediate immune response (CMI), and challenge results were used to evaluate the immune efficacy of the vaccine candidates. Pt-Dd induced the highest level of enzyme-linked immunosorbent assay antibodies and high levels of CMI, showing a significantly (p < 0.05) high level of interferon γ, interleukin-4, and major histocompatibility complex II expressions in peripheral blood mononuclear cells at 48 h post-infection. The challenge results showed that Pt-Dd, inactivated FAdV-4 vaccine, as well as fiber-1 induced the best protection (100%), followed by fiber-2 (80%) and penton (67%). The present study showed that FAdV-4-Pt-Dd and recombinant fiber-1 are good FAdV-4 vaccine candidates and could be used to replace the tissue-sourced inactivated FAdV-4 vaccine.

**Importance:** FAdV-4-Pt-Dds were discovered in FAdV-4 infected cells, and which were also assembled in cells transduced with recombinant human adenovirus expressing fiber-1, fiber-2, and penton base. FAdV-4-Pt-Dds internalize into cells with high efficiency, so that it can be used for delivery vaccine antigen or drugs. Immunization chickens with Pt-Dd and fiber-1 obtained by transduction HEK-293T cells induced significant high level humoral and cellular mediated immune responses, and also 100% challenge protection in chickens indicating that they are good FAdV-4 vaccine candidates. What more, the Pt-Dd obtained by transduction HEK-293T cell would have no DNA and adenovirus contamination as adenovirus could not package in HEK-293T cells.

## 1. Introduction

Adenoviruses (Ad) are large, non-enveloped viruses with an icosahedral nucleocapsid containing a double-stranded DNA genome of approximately 36 KD (1). They have a broad range of vertebrate hosts and, in humans, more than 50 distinct adenoviral serotypes have been found to cause a wide range of illnesses, from mild respiratory infections in children (known as the common cold) to life-threatening multi-organ diseases in people with a weakened immune system (2). The capsid of the Ad is composed of three major capsid proteins (hexon, penton base, and fiber) and four minor capsid proteins (IIIa, VI, VIII, and IX; also known as cement proteins) (3). The pseudo-T=25 icosahedral capsid has 12 pentons, each with trimeric fiber proteins and 240 trimeric hexons (4).

In some serotypes of Ad, during their replication cycle in the infected cells the penton base proteins and the attached trimeric fiber assemble into dodecameric virus-like particles called Ad penton-dodecahedron (Pt-Dd) [14]. Both the penton base and trimeric fiber are responsible for Ad attachment and endocytosis (5, 6). Native VLPs can be observed in Ad3-, Ad4-, Ad9-, Ad11-, and Ad15-infected cells (7–9). It can also be assembled spontaneously in the baculoviral system (6, 10) and *Escherichia coli* (11) upon expression of the penton and fiber gene of Ad3. No such particles have been found for Ad2 or Ad5 (12–14). This particle shows a remarkable cell penetration ability with 200–300 thousand VLPs per cultured cell (15, 16); thus, it can be engineered to deliver several millions of foreign cargo molecules into a single target cell (15–18).

Pt-Dd has been exploited to deliver antigen proteins (19, 20) or small molecule drugs (21) linked by an additional domain to the penton base protein. In a previous study, Pt-Dd was used as a cancer vaccine vehicle to deliver model antigen ovalbumin (OVA) fused with the linker WW domain from the ubiquitin ligase Nedd4 (20). Pt-Dd can efficiently deliver WW-OVA in vitro and the Pt-Dd/WW complex can be readily internalized by dendritic cells. Immunization with WW-OVA/Pt-Dd induced OVA-specific CD8 + T cells immune response and robust humoral responses in mice resulting in 90% protection against B16-OVA melanoma implantation in syngeneic mice (20). Pt-Dd was also used to develop multivalent vaccination-carrying influenza epitopes. With this platform, the immunodominant epitopes of the influenza M1 were properly presented by human dendritic cells, thus triggering the efficient activation of antigen-specific T cell responses and inducing cellular immunity in vivo in chickens (19).

Fowl adenoviruses (FAdVs) are classified into five species (A–E) (22) and twelve serotypes (FAdV 1–7, 8a, 8b, 9–11) (23). FAdV-4 is the etiology of the hydropericardium hepatitis syndrome (HHS) that is characterized by hydropericardium, hepatitis, and nephritis (23–25). FAdV-4 is the predominant serotype of FAdVs in southwestern China. As reported, about 86.4% (19/22) isolates were FAdV-4 (26). FAdVs have been found to infect 2- to 5-week-old neonatal chicks (26), causing a mortality of approximately 30% to 70% (27, 28). Inactivated vaccines (29–31), attenuated live vaccines (32, 33), and recombinant vaccines have been used to control HHS and these types of vaccines have been proven to be effective in protecting against FAdV infection. In a previous study, scientists purified Pt-Dd produced by FAdV-8b via density gradient centrifugation, which induced neutralizing antibodies and cytotoxic T-cell responses in breeders, as well as successfully prevented clinical disease in progeny via maternal antibody transfer (34). FAdV-4 might also produce Pt-Dd during infection; therefore, the present study utilized the human Ad5 to produce recombinant FAdV–4–Pt-Dd and used this as a vaccine to control the FAdV-4 infection.

## 2. Results

### 2.1. Construction of recombinant viruses

Shuttle vectors containing fiber-1, fiber-2, and penton base of FAdV-4 under the control of the cytomegalovirus immediate early (CMV) promoter were constructed and verified by sequencing (data not shown). By recombination with Ad backbone vector pAdEasy-1 in BJ5183 cells, recombinant adenoviral plasmids, pAd-fiber-1, pAd-fiber-2, and pAd-penton base were obtained. Then, the recombinant plasmids were linearized with endonuclease *pac* I and transfected into HEK293A to generate recombinant Ad (rAd), rAd-f1 (expression fiber-1), rAd-f2 (expression fiber-2), and rAd-p (expression penton base). After approximately 10 days incubation, the rAds were successfully packaged with characteristic cytopathic effect into transfected cells, while the mock-transfected cells (control samples) retained their singularity (Fig. 1A).

**Fig. 1.**
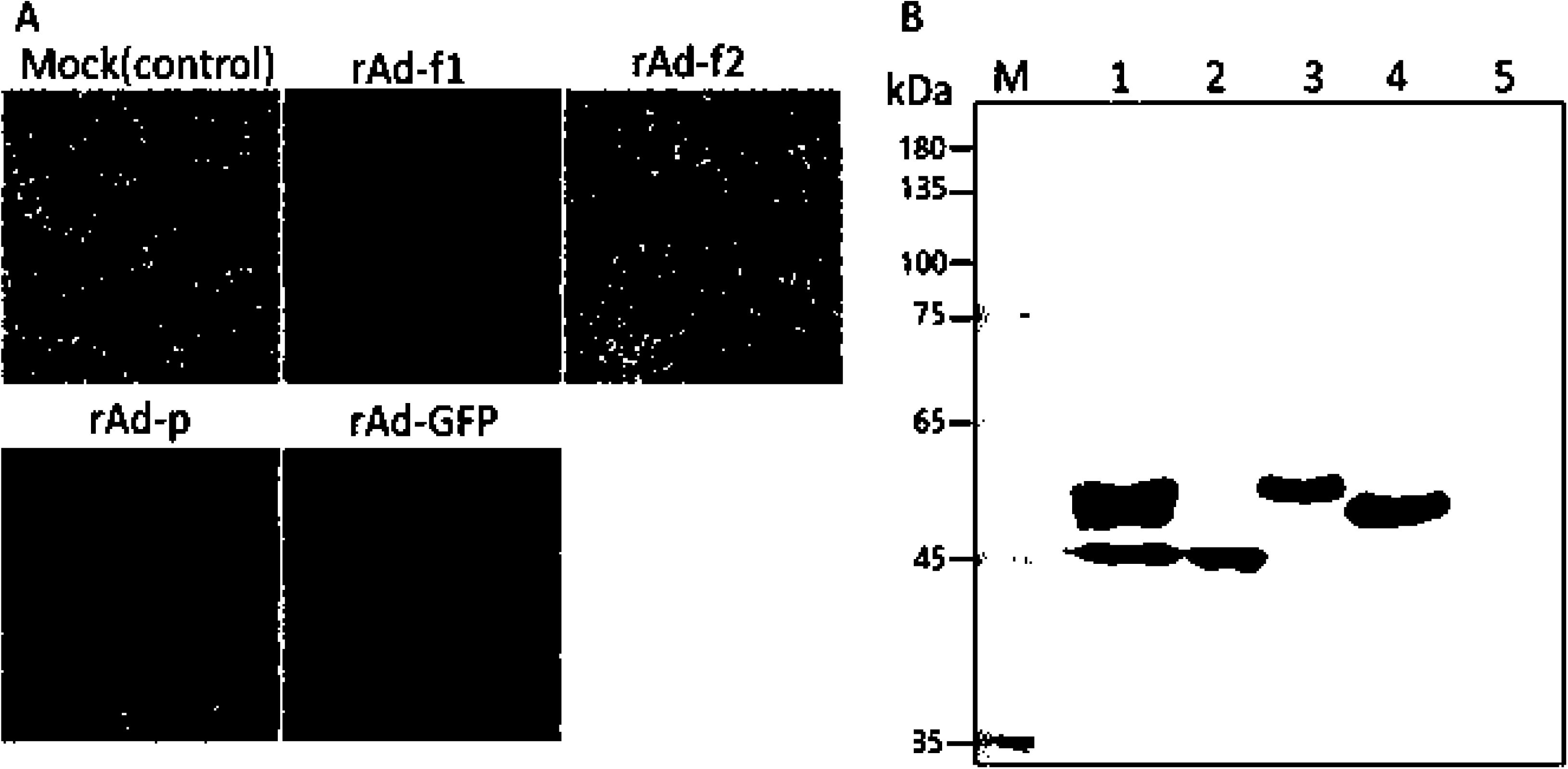
Package rAd in HEK293A cells and identification of recombinant protein expressions by western blotting. (A) Micrographs of HEK293A cells at 10 days post-transfection with non-plasmid (mock) or linearized pAd-fiber-1, pAd-fiber-2, and pAd-penton base. rAd-GFP was used as the control (2 days post-viral inoculation); (B) Western blotting identification fiber-1, fiber-2, and penton base protein expression in transfected HEK293A cells with polyclone antibodies against fiber-1, fiber-2, or penton base of FAdV-4, respectively. Line 1 was the positive control added with the cell lysis from chicken embryo fibroblast infected with FAdV-4 and Line 5 was added with the proteins from cells infected with rAd-GFP. Lines 2, 3, and 4 corresponded to fiber-1, fiber-2, and penton base, respectively.

The obtained rAds, rAd-f1, rAd-f2, and rAd-p, were purified three times using the plaque-purified method and titered in HEK293A cells. The titers of these three rAd were all 4.1 × 10^8^ vp mL^−1^. The expressions of the proteins, fiber-1, fiber-2, and penton base with predicated sizes of 48.5, 53.8, and 58.8 kDa were confirmed by western blotting. As shown in Figure 1B, the protein bands corresponding to fiber-1, fiber-2, and penton base are found in Line 2, 3, and 4, respectively, in which the proteins derived from cells infected with rAd-f1, rAd-f2, and rAd-p were separated. Three protein bands were observed in Line 1 running proteins from LMH cells (Chicken Liver Hepatocellular carcinoma cell line) infected with FAdV-4. No band was observed in Line 5, in which the proteins from cells infected with rAd-GFP (rAd expression green fluorescent protein) were added.

### 2.2. Fiber-1, fiber-2, and penton base derived from FAdV-4 or rAds formed Pt-Dd

In the cell lysis derived from LMH infected with FAdV-4, the virions of FAdV-4 approximately 100 nm in size and the Pt-Dd approximately 50 nm in size were observed by TEM. The morphology of the virions of FAdV-4 was the classic dodecahedron symmetry Ad structure and the Pt-Dds showed a similar highly ordered architecture of dodecahedrons.

To check whether Pt-Dd could also assemble by the expressed recombinant proteins, HEK293A cells, as well as HEK293T cells, were infected with rAd-f1, rAd-f2, and/or rAd-p. Pt-Dds became assembled and were observed in HEK293A cells and HEK293T cells co-infected with rAd-f1, rAd-f2, and rAd-p (Fig. 2B and 2C). In addition, the Ad5 virions were found in infected HEK293A cells (Fig. 2B), but not in HEK293T cells. The penton base alone also formed dodecahedrons in HEK293A cells infected with rAd-p (Fig. 2D), but not in cells infected with rAd-f1 and rAd-f2 (Fig. 2B or Fig. 2C).

**Fig. 2.**
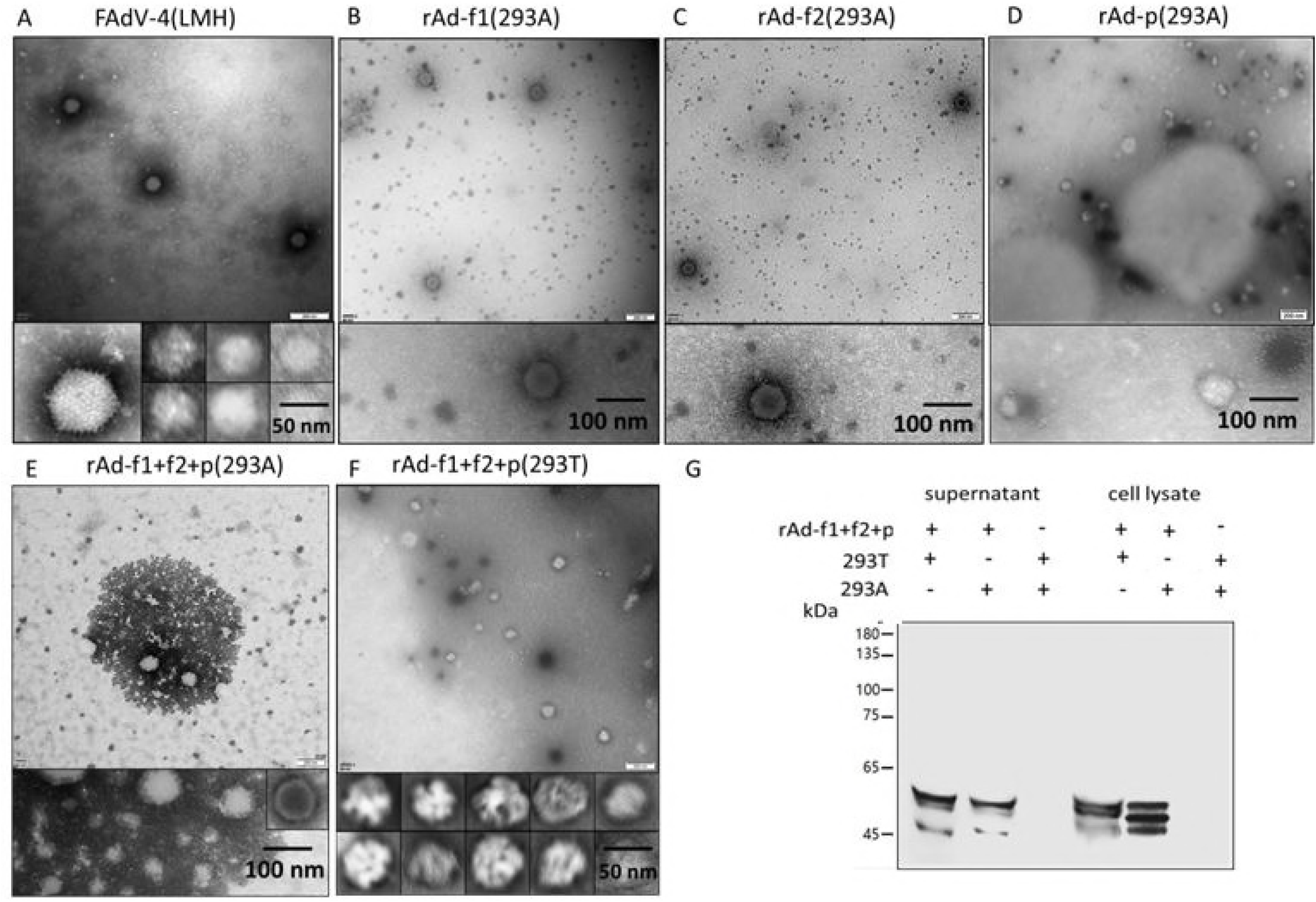
Transmission electron microscopy micrographs of Ad viral particles and/or Ad viral-like particles in cell culture media. (A) Viral particles in cell culture media of LMH infected with FAdV-4; (B, C, D) Viral particles in cell culture media of HEK293A cells infected with rAd-f1, rAd-f2, or rAd-p, respectively; (E) Viral particles in cell culture media of HEK293A cells co-infected with rAd-f1, rAd-f2, and rAd-p; (F) Viral particles in cell culture media of HEK293T cells co-infected with rAd-f1, rAd-f2, and rAd-p; (G) Western blotting detection of fiber-1, fiber-2, and penton base expression in supernatant and cell lysis of HEK293A or HEK293T cells co infected with rAd-f1, rAd-f2, and rAd-p.

As Ad5 virions would not assemble in non-complementing HEK293T cells and Ad5 has no Pt-Dd produce, the obtained Pt-Dd in Figure 2F could only be derived from the overexpressed fiber-1, fiber-2, and penton base of FAdV-4. In addition, Pt-Dd in cell culture supernatants and cell lysates were detected with western blotting (Fig. 2E) and the concentration of the Pt-Dd in cell supernatants was approximately 50 µg/mL.

### 2.3 Internalization of FAdV-4 VLPs

To generate Pt-Dd without the influence of human Ad virions, rAd-f1 and rAd-f2 together with rAd-p were used to infect HEK293T cells. Culture media were collected at 48 h post-infection. The cell culture media from HEK293T cells infected with rAd-GFP were used as the control.

The internalization of Pt-Dds into HeLa cells was analyzed by observation under a confocal microscopy after staining with polyclone antibodies against fiber-1 and Cy3-labeled secondary antibodies. Pt-Dd was observed to gather around cells as early as 10 min after inoculation (Fig. 3A) and the internalization was finished within 2 h as almost all the fluorescence was in the cells by that time (Fig. 3E). The internalization process was clearly related to the time of incubation (Fig. 3). Each cell that had been incubated with the collected supernatant converged with thousands of Pt-Dds. Fiber-1 alone and GFP could not be internalized into cells as they did by Pt-Dds (data not shown).

**Fig. 3.**
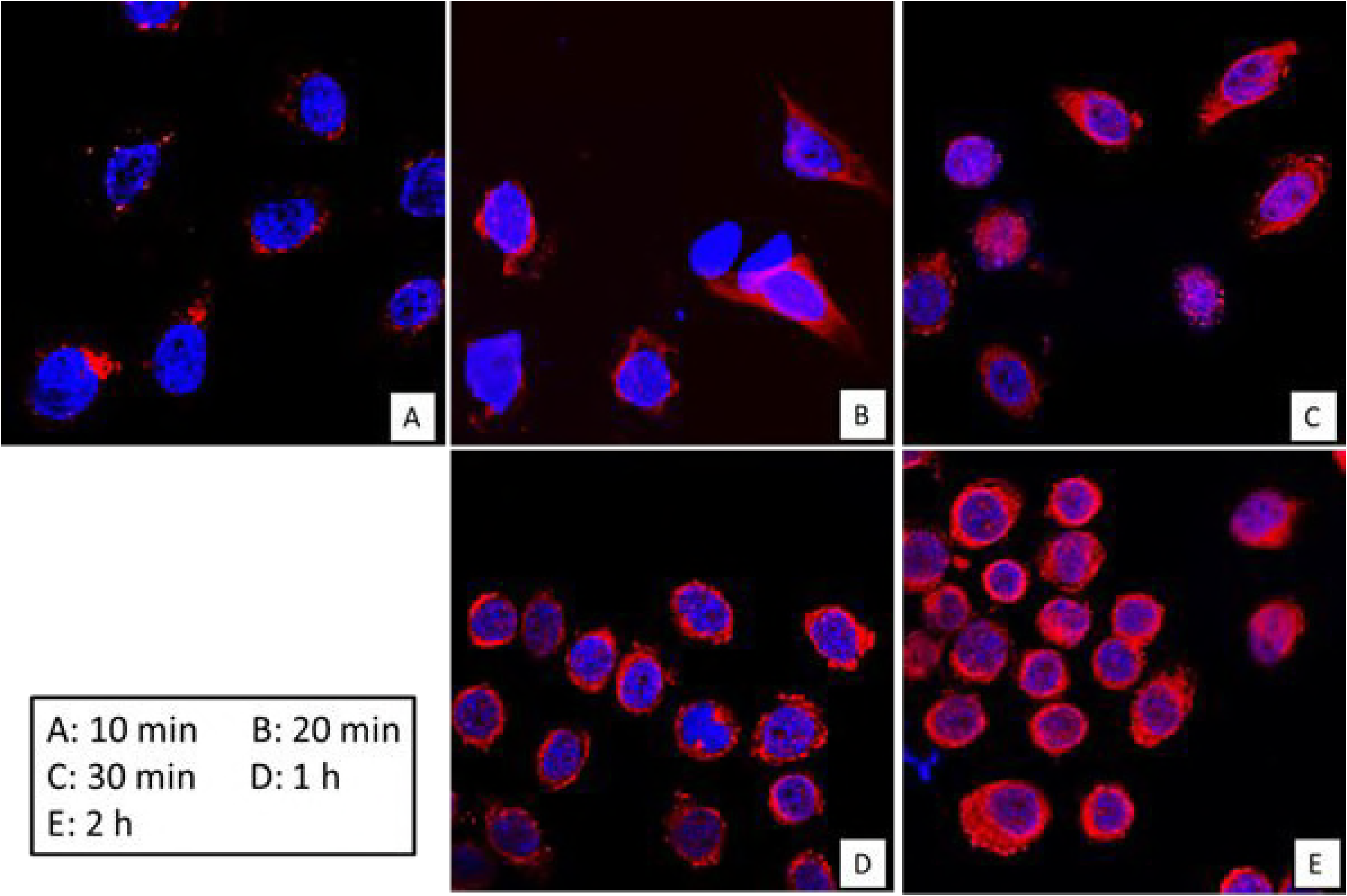
FAdV-4–Pt-Dd internalized into HeLa cells as determined by confocal microscopy. FAdV-4– Pt-Dd are shown in red with a Cy3-conjugated mouse antibody against fiber-1, while the nuclei are counterstained in blue with Hoechst 33342.

### 2.4 Humoral and cellular immune responses against FAdV-4 following chicken vaccination

Chickens were immunized on day 7 with approximately 5 µg (0.1 mL) of recombinant proteins obtained by transduction of HEK293T cells with the corresponding rAd and the challenged chickens were administered with FAdV-4 on day 21. The chicken sera were collected on days 14, 21, and 35 post-immunization and were detected by enzyme-linked immunosorbent assay (ELISA) with the mixed proteins (*E. coli* expressed fiber-1, fiber-2, and penton base; 0.2 µg each in one well) as reported previously (35). The results showed that the antibodies reached a peak on day 28, i.e. 7 days post-challenge and then decreased slightly. On day 21, the average antibody titers (log10) in chickens immunized with Pt-Dd, fiber-1, fiber-2, penton, or inactive vaccine were 4.56 ± 0.67, 3.54 ± 0.20, 3.82 ± 0.21, 3.82 ± 0.21, and 3.82 ± 0.00, and the chickens immunized with Pt-Dd gained significantly higher (p<0.05) levels of antibodies than that of the other immunized groups (Fig. 4D).

**Fig. 4.**
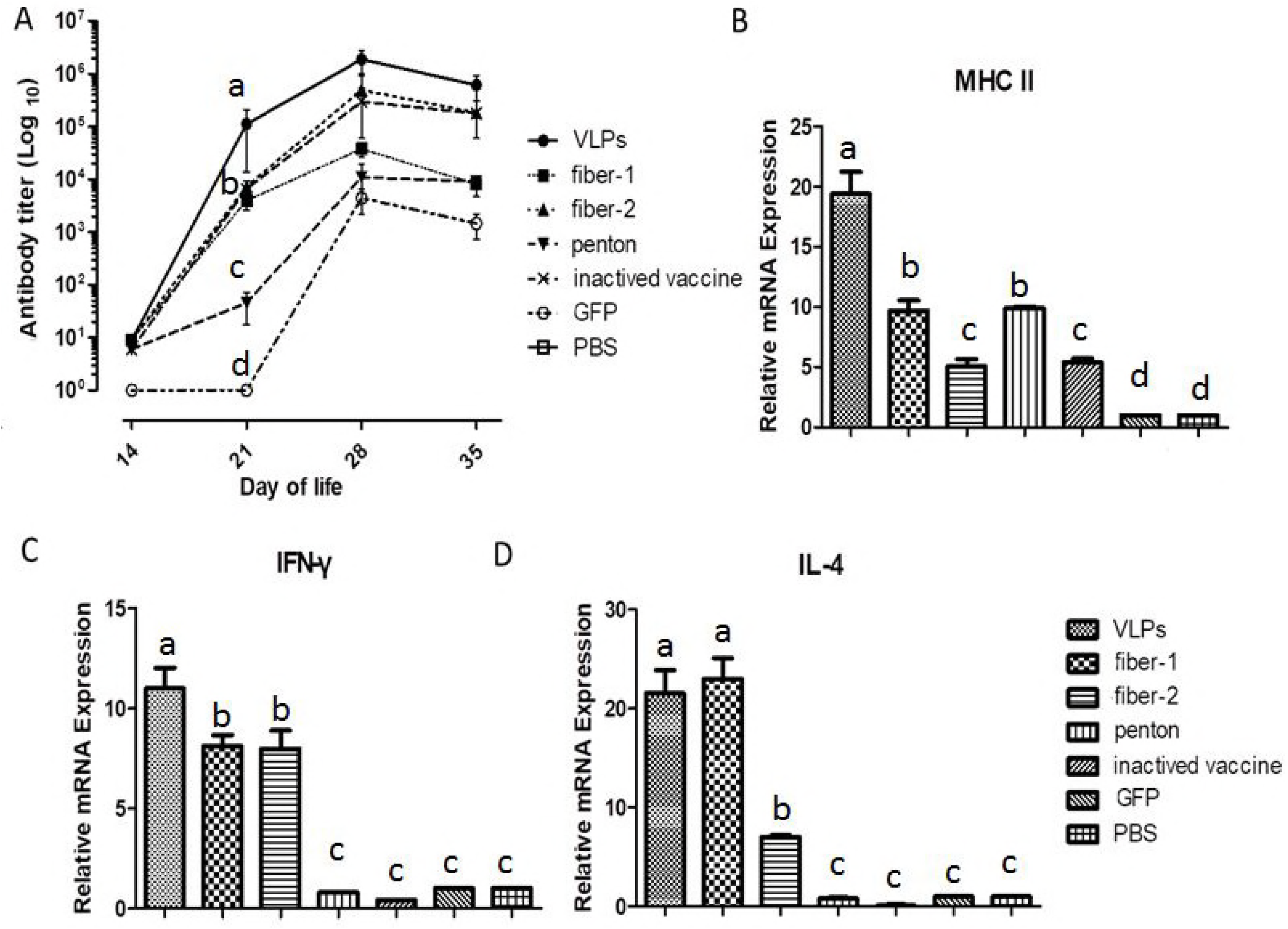
Humoral and CMI response induced by immunization. (A) Detection FAdV-4 antibodies with ELSIA; (B, C, D) Detection of CMI related molecule expressions in PBMC by RT-qPCR.

Cellular immune responses elicited by vaccination were evaluated by qualification of interferon γ (IFN-γ), interleukin-4 (IL-4), and major histocompatibility complex II (MHC-II) expression using real time quantitate Polymerase Chain Reaction (RT-qPCR) as reported previously(36). The peripheral blood mononuclear cells (PBMCs) were collected at 24 h post-challenge and the expressions of IFN-γ, IL-4, and MHC-II were detected. As shown in Figure 4, vaccination with Pt-Dd significantly increased the expressions of IFN-γ, IL-4, and MHC-II in PBMC (Fig. 4 A, B, C) compared with those given fiber-2 or penton base. Fiber-1 also induced high levels of IL-4 expression.

### 2.5 Challenge protection test

All chickens except those in the non-infection control group were challenged with 10^5.5^ TCID_50_ of FAdV-4 SX15. The FAdV-4 SX15 challenge resulted in 73.3% (11/15) mortality in the challenge control group. Immunization with Pt-Dd, fiber-1, fiber-2, penton base or inactivate vaccine reduced the mortality from 73.3% to 0%, 0%, 20%, 33.3%, and 0%, respectively, and the corresponding survival ratios were 26.7%, 100%, 80%, 67.7%, and 100%, respectively (Fig. 5).

**Fig. 5.**
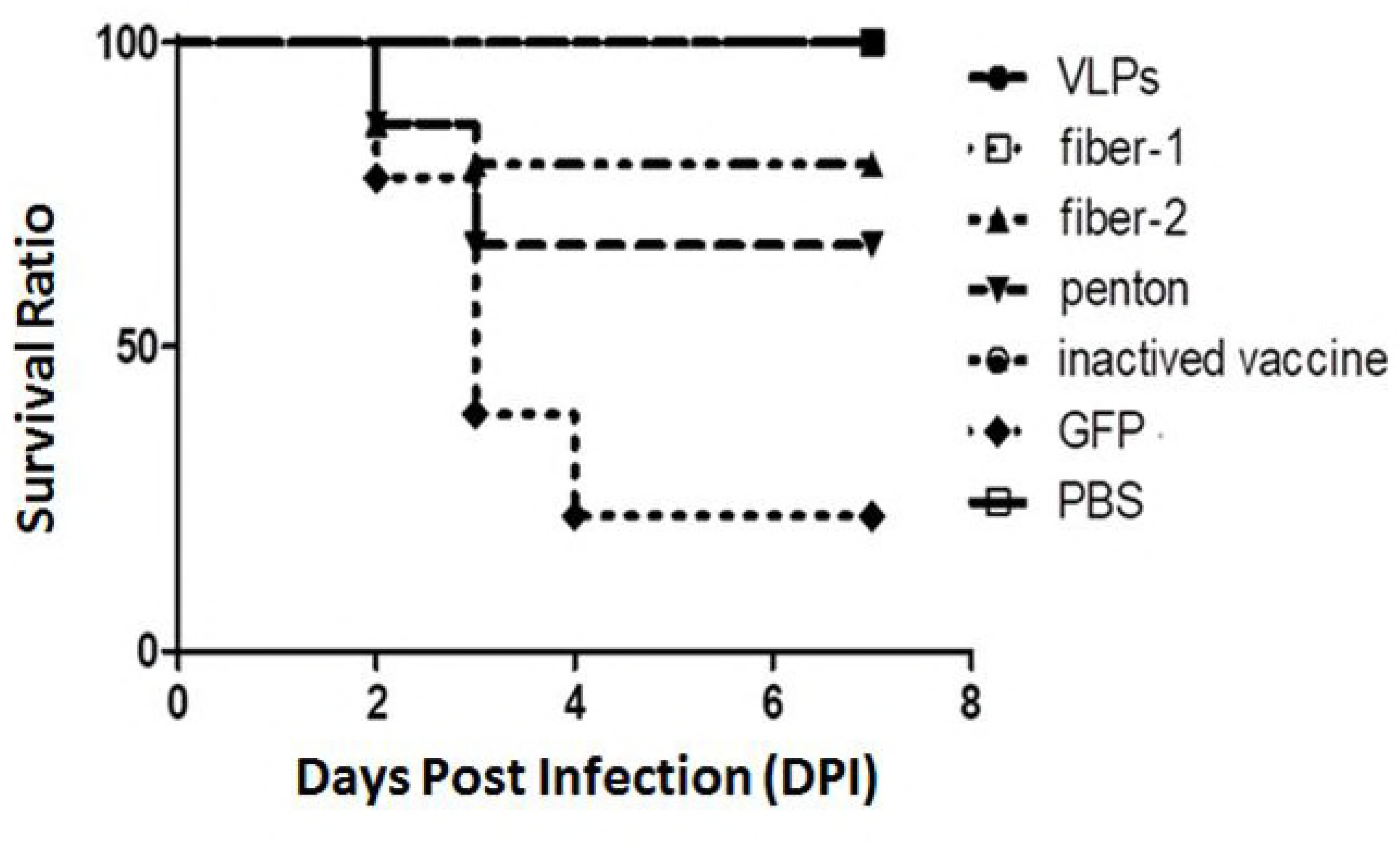
Survival ratios of chickens to the challenge of virulent FAdV-4. Chickens were challenged with 10^5.5^ TCID_50_ of FAdV-4 SX15 strain and the survival ratios were recorded for 7 days when all challenged chickens were recovered.

### 3.6 Histology and immunohistochemistry

Histology and immunohistochemistry analyses were used to distinguish the micro-level differences among the immunized groups because the proteins derived from the Ad provided complete or almost complete protection to the chickens. The most serious micro lesions were observed in the livers of chickens in the control group given GFP. The hepatic lobule structures were completely missing. The hepatocyte tubes were arranged haphazardly with severe congestion and inflammatory cell infiltration. Liver cells appeared necrotic with nuclear fragmentation. Immunization with penton reduced the pathological changes mentioned above (Fig. 6). However, the hepatic lobule structures were still not very clear and parts of the liver cells appeared necrotic with nuclear fragmentation (Fig. 6). Fiber-2 also did not protect liver microstructures well; however, the lesions were only slight and necrotic liver cells were even fewer (Fig 6). Comparatively, Pt-Dd, fiber-1, and the inactive vaccine immune group provided better protection against FAdV-4 infection based on observations of the pathological tissue slices. The hepatic lobules had a clear structure and the hepatocyte tubes were arranged orderly and closely in the livers from these groups. Meanwhile, ecchymosis in the hepatic lobules was observed in the livers from both groups, and immunohistochemical analysis indicated that serious infection occurred in chickens injected with GFP and penton. Fewer FAdV-4-positive signals were detected in the liver sections of chickens from groups immunized with FAdV-4 VLPs and the inactive vaccine immune group, although the difference was not significant among the VLP, fiber-1, fiber-2, and inactive vaccine immune groups (Fig 7).

**Fig. 6.**
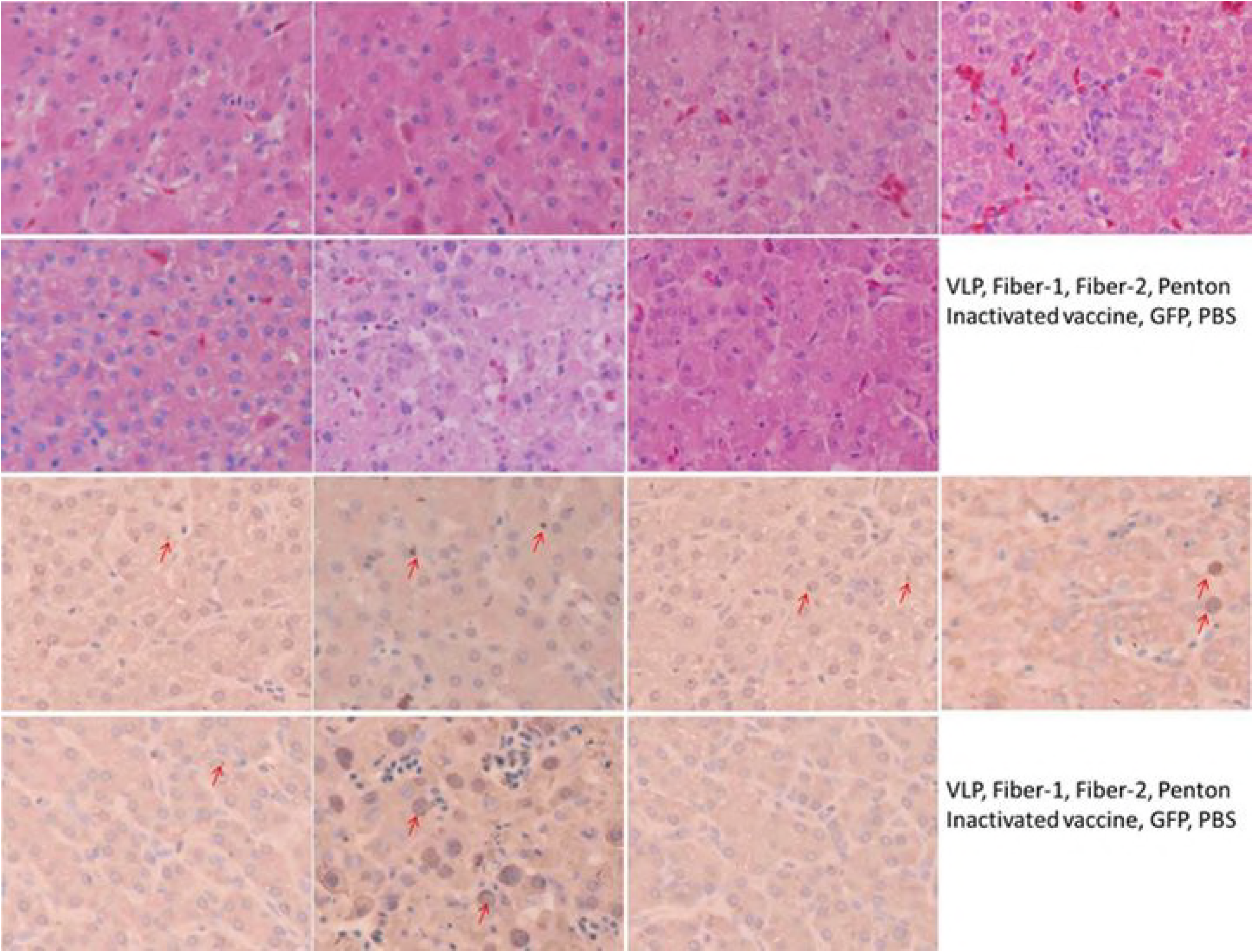
Observation of micro-lesions and detection of viral antigens in the liver with hematoxylin and eosin staining and histology and immunohistochemistry analyses. (A) Histological analysis of the liver (hematoxylin and eosin). The lesions are shown as cell body boggy, structure blurring, nuclear boggy, even shattering, melting and so on. The most clear lesions are observed in the tissues from the non-immunized chickens and then from the penton and fiber-2 groups; (B) Detection of viral antigens using histology and immunohistochemistry analyses. Viral positive signals (dark brown) were detected in cells or hepatic sinusoid (highlighted by the red arrows)

**Fig. 7.**
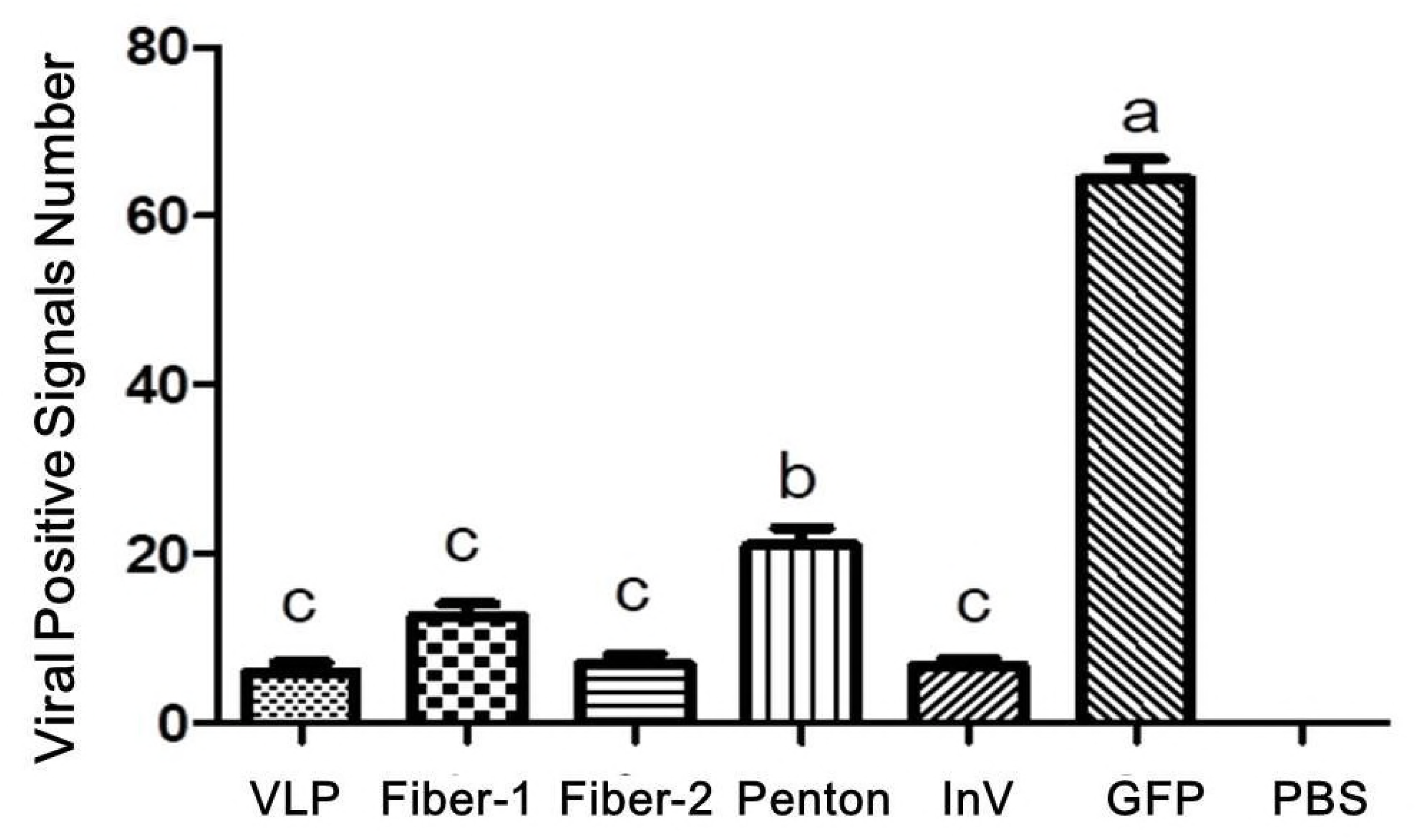
Calculated viral positive signals in liver sections. All data were collected from three slices and five random selected views. Data are presented as mean ± standard deviation and significant differences are marked with different letters (p ≤ 0.05).

## Discussion

Ad vectors are the most commonly applied viral vectors for gene therapy and delivery of vaccine antigens. The Ad vectors that are used as vaccines are mostly replication-defective with certain essential viral genes deleted and replaced by a foreign gene expression cassette(37). In the present study, replication-defective Ad5 was used with expression fibers and penton base of FAdV-4. The Ad5 vector has essential genes, E1A and E1B, deleted and replaced by a cassette of expression foreign genes under the control of the CMV promoter. The E1A code immediate early proteins, which play a role in the initiation of the expression of the ∼20 delayed early genes in the E1B, E2, E3, and E4 transcription units that are essential for Ad replication. These proteins are induced by E1A and facilitate adenoviral replication by altering the expression of a multitude of cellular genes(37). The E1B proteins, in general, inhibit the cell apoptosis caused by adenoviral infection. These deleted Ad5 vectors are constructed from plasmids or Ad DNA containing the genetically modified Ad genome, and the vectors are grown up on complementing cell lines such as HEK293A, PER.C6, or N52.E6 expression of the E1A and E1B genes(38). Thus, these deleted Ad5 vectors would undertake continuous passage in HEK293A. In HEK293T cells without E1A and E1B genes, Ad5 would not complete packing, with the carrying foreign genes expressed at high levels because they are controlled by the CMV promoter. In the present study, Ad5 virions, as well as Pt-Dd of FAdV-4 (Ad5 has no Pt-Dd) were observed in HEK293A cells that had been infected by rAd-f1, rAd-f2, and rAd-p; however, no Ad5 virions, only Pt-Dds, were found in HEK293T infected cells. Accordingly, it would be convenient to obtain ectogenic Pt-Dds without contamination with Ad virions by transduction of non-complementing cells with the Ad5 vector.

Dodecahedra composed of 12 pentons are found in cells infected with Ad3(39). These are synthesized in abundance with 5.5 × 10 ^6^ Dds produced per one infectious virus(40). As calculated, approximately 200-fold more Dds than total viral particles are found in Ad3-infected cells, at 24 h.p.i. Ad Dd is different to other known VLPs of non-enveloped viruses, such as human papilloma virus or Norwalk VLPs, whose VLPs morphologically mimic native virions. Ad Dd is smaller and lacks several structural components of the Ad viral partial resulting in functional and structural properties that are different from those of the virus capsid. For example, Dd is able to attach to and penetrate cells via interaction with heparan sulfate, a pathway not used by the virus of origin, Ad3(41, 42). Owing to its high endocytosis capacity, Pt-Dd has been investigated extensively as an alternative vector for antigen protein(43, 44) or small molecule drugs(45). In the present study, the characteristics of Pt-Dd from fAdV-4 were studied. By co-infection of HEK293A or HEK293T cells with recombinant Ad5 expression fiber-1, fiber-2, and penton of fAdV-4, Pt-Dd were properly packaged, and they could be efficiently internalized into human cells (Fig. 3) and chicken embryo fibroblast (data not shown). Because fAdV-4 is not from a human source, there is less chance that related antibodies would be found in human; therefore, the fAdV-4 derived Pt-Dd could be a good drug delivery system for human disease therapy.

Ad3 Dds exhibit remarkable stability and can be stored for long periods at 4°C and even at room temperature(46). Thawing and reconstitution with water after being frozen and lyophilized did not significantly influence their integrity in the present study. As previously reported, Dds retain their particulate integrity in human serum at 37°C for at least 2 h(46). Accordingly, Dds can be conveniently stored and transported, and can be used as a vaccine or drug delivery system under various climates. Although we did not check the stability of Pt-Dd of FAdV-4, they can penetrate HeLa cells after 4 weeks of storage at 4°C (data not shown).

In the present study, Pt-Dd of FAdV-4 VLPs were produced using Ad5, and the immune response induced by Pt-Dd were evaluated. Pt-Dd is readily internalized by dendritic cells and is one type of principal antigen-presenting cells for T cells. Pt-Dd, as well as their carrying antigens, might induce strong cell mediated immune response. The immunization of OVA linked to Ad3 Pt-Dd by the WW domain developed OVA-specific CD8 + T cells and robust humoral responses in mice. Proteins or peptides alone do not easily illicit an immune response. Pt-Dd, by improving the antigen delivery, played the role of an adjuvant. In the present study, the activated cells mediated immune responses by Pt-Dd were also detected and presented as significant higher levels of IFN-γ, IL-4 and MHC-II expressions in PBMC after boost immunization. In addition, Pt-Dd also induced higher levels of humoral immune response with significant high levels of antibodies in serum detected by ELISA. In a previous study, the purified FAdV-8b VLPs also induced neutralizing antibodies and cytotoxic T-cell responses in breeders as well as successfully preventing clinical disease in progeny via maternal antibodies transfer (47). Accordingly, the Pt-Dd of FAdVs might be an ideal vaccine candidate for the control the related diseases.

In the present study, fiber-1, but not fiber-2, provided relatively better protection to the FAdV-4 challenge. This result is different to our previous study and several other related studies, in which fiber-2 induced better protection against FAdV-4 infection than fiber-1 (35, 48). These differences might be related to the protein structure, with their actual structure shown in eukaryocytes, but not in *E. coli* or other expression systems. Further studies are required for to reveal the reason behind these differences.

In summary, by using defective Ad5, we successfully packaged Pt-Dd of FAdV-4. The obtained Pt-Dd could penetrate different types of cells with high efficiency indicating that it could be a new drug or vaccine antigen deliverer. Immunization with Pt-Dd induced high levels of cell mediate and humoral immune response in chickens and offered complete protection with 100% survival.

## Conflict of Interest

The authors declare no conflict of interest.

## Acknowledgements

This work was supported by the National Natural Science Foundation of China (No. 31672581).

## 3 Materials and methods

### 3.1 Viruses and cells

The human type 5 replication-defective Ad5 was purchased from TaKaRa (Dalian, China) and rAd were propagated and titered in HEK-293A cells. All cells were grown in Dulbecco’s modified Eagle’s medium (DMEM) and supplemented with 10% fetal bovine serum (FBS), 2 mM L-glutamine, 100 U/mL penicillin, and 100 µg/mL streptomycin at 37°C in a humidified atmosphere of 5% CO_2_. The fowl adenovirus strain SX15 was stored and propagated by our laboratory.

### 3.2 Construction of rAds (rAd-f1, rAd-f2, and rAd-p)

The respective open reading frames of fiber-1, fiber-2, and penton base were amplified by polymerase chain reaction (PCR) from viral DNA (FAdV-4 isolate SXD15, refer to KU569296.1) using the primers listed in Table 1. The PCR amplicons of fiber-1, fiber-2, and penton base were cloned into a pAd-shuttle-CMV vector. The recombinant adenoviral vectors were generated by homologous recombination of linearized transfer vectors with pAdEasy-1 in *E. coli* BJ5183 and confirmed by restriction enzyme digestion (New England Biolabs)(49). The rAds were generated by transfection of 1 µg plasmids (PacI linearized) using 3 µL of Trans Fast TM Transfection Reagent (Promega, Madison, USA). When 90% of the cells showed cytopathic effect, Ads were released by three cycles of rapid freezing and thawing and stored at –80°C after the addition of 10% glycerol.

**Table 1.**
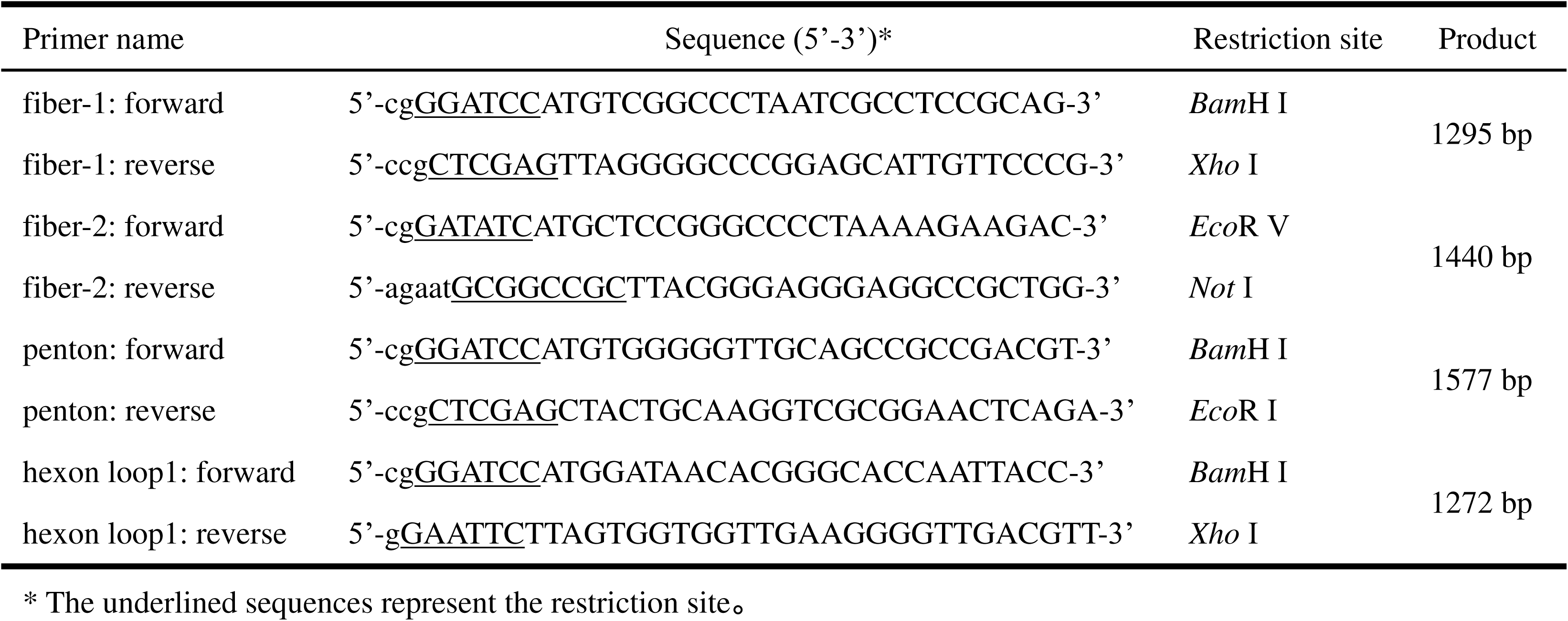
Primers used for recombinant adenoviruses construction.

### 3.3 Western blot

The cell lysates were separated by 10% sodium dodecyl sulfate polyacrylamide gel electrophoresis (SDS-PAGE) and transferred onto a nitrocellulose membrane (Pall Corporation). The 293A cells infected with rAd-GFP were used as the negative control. The membrane was incubated overnight in blocking solution (10% fat-free milk in phosphate buffer solution (PBS), PBS-M) at room temperature and incubated with FAdV-4 fiber-1, fiber-2, and penton base specific mouse antiserum for 2 h. The membrane was subsequently reacted for 1 h with goat anti-mouse IgG conjugated with horseradish peroxidase (AS003) at a dilution of 1/2000 in PBS-M. Detection was performed using chemiluminescence luminol reagents (Super Signal West PicoTrial Kit, Pierce). Utilizing gray scan and comparison with the known concentration marker bands, the concentrations of proteins were calculated based on the obtained gray value of the bands.

### 3.4 Observation Ad virion and Pt-Dd by electron microscopy

Virus samples were applied to the clean side of carbon on mica (carbon/mica interface) and negatively stained with 1% sodium silicotungstate, pH 7.0. Micrographs were taken with a transmission electron microscopy (TECNAI G2 SPIRIT BIO), under low-dose conditions with the microscope (FEI, America) at 100 kV and a maximum magnification of 300,000.

### 3.5 Observation Pt-Dd internalization into cells by confocal microscopy

HeLa cells were grown overnight on glass coverslips (approximately 10^5^ cell/cm^2^) in DMEM supplemented with 10% FBS at 37°C under 5% CO_2_ atmosphere. Cells were incubated for different periods with a 0.6 nM concentration of the FAdV-4 VLPs. After a given period of entry, cells were washed with PBS, fixed, and permeabilized with cold methanol for 10 min. Cells were then incubated with anti-fiber-1 mouse polyclonal serum at 1/200 in 50 µL 3% bull serum albumin and subsequently detected in red by Cy3-conjugated goat anti-mouse antibody (BA1035, Bosterbio, Wuhan) that was diluted 1/200 in the same buffer. Cell nuclei were counterstained in purple with propidium iodide (5 µg/mL). Laser scanning confocal microscopy was performed on an MRC600 (Andorra, England).

### 3.6 Immunization and challenge

Day-old Specific Pathogen Free chickens were randomly divided into 7 groups of 15. The chickens in the first 5 groups received 0.5 µg of FAdV-4 Pt-Dd VLPs, fiber-1, fiber-2, penton base, and inactivated vaccine of FAdV-4 (inactive vaccine) by intramuscular injection on days 7 and 14. The other two groups received PBS.

At day 21, all chickens except those in the negative control group (PBS), were challenged with 10^5.5^ TCID_50_ of FAdV-4 SX15. Sera (n = 3 per group) were collected on days 14, 21, 28, and 35 to determine serum antibodies. Additionally, blood was collected at 24 h post-challenge to measure cell mediate immune response (CMI) related gene expression in PBMC.

### 3.7 Histopathology and immunohistochemistry analyses

Three surviving chickens from each group were randomly selected and euthanized by CO_2_ asphyxiation at 6-days post-challenge. Liver samples were fixed in 10% neutral formalin for 52 h at room temperature. After routine treatment, liver tissues were embedded in paraffin wax and cut into 4 to 5 µm slices. For micro lesion observation, the slices were stained with hematoxylin and eosin. In addition, immunohistochemical staining was conducted to detect viral antigens in the livers. Briefly, slices were incubated with homemade mouse anti-FAdV poly antibody (diluted 1:200) for 1 h and then reacted with horseradish peroxidase-conjugated rabbit anti-mouse IgG (Solarbio, Beijing, China) for 1 h. DAB (3,3-diaminobezidin) was used as the chromogenic substrate. After counterstaining with hematoxylin and eosin, the slides were dehydrated and mounted. Slides without the primary antibody were used as negative controls.

### 3.8 Enzyme-linked immunosorbent assay (ELISA)

ELISA plates (Guangzhou Jet Bio-Filtration Co., Ltd., China) were coated with 0.2 µg (each)/well of mixed purified fiber-1, fiber-2, and penton (expressed in *E. coli* and stored by our laboratory). Then, the plates were reacted with sera from chickens immunized with the corresponding proteins. Commercial peroxidase-conjugated rabbit anti-chicken IgG (Sigma) was used as the secondary antibody. Following incubation with tetra-methylbenzidine substrate (Tiangen Biotech (Beijing) Co., ltd., China), the reaction was stopped with 0.5 M sulfuric acid and the optical density of each well was measured at a wavelength of 450 nm.

### 3.9 Statistical analysis

One-way ANOVA, followed by a Tukey post-test, was performed for multiple comparisons. Differences were considered significant at p≤0.05. Statistical analysis was run using SPSS (version 22) software.

## References

1. Davison AJ, Benkő M, Harrach B. 2003. Genetic content and evolution of adenoviruses. Journal of General Virology 84:2895–2908.

2. Tsoline K, Phyllis F, S. HM. 2003. The impact of adenovirus infection on the immunocompromised host. Reviews in Medical Virology 13:155–171.

3. San Martín C. 2012. Latest Insights on Adenovirus Structure and Assembly. Viruses 4:847–877.

4. Zubieta C, Schoehn G, Chroboczek J, Cusack S. 2005. The Structure of the Human Adenovirus 2 Penton. Molecular Cell 17:121–135.

5. Wickham TJ, Mathias P, Cheresh DA, Nemerow GR. 1993. Integrins alpha v beta 3 and alpha v beta 5 promote adenovirus internalization but not virus attachment. Cell 73:309–319.

6. Szolajska E, Burmeister WP, Zochowska M, Nerlo B, Andreev I, Schoehn G, Andrieu JP, Fender P, Naskalska A, Zubieta C. 2012. The structural basis for the integrity of adenovirus Ad3 dodecahedron. Plos One 7:e46075.

7. Norrby E. 1966. The Relationship between the Soluble Antigens and the Virion of Adenovirus Type 3. 1.Morphological Characteristics. Virology 28:236–248.

8. H G, H B, H F, R W. 1967. The structure of group II adenoviruses. Journal of General Virology 1:553–560.

9. Norrby E, Skaaret P. 1968. Comparison between soluble components of adenovirus types 3 and 16 and of the intermediate strain 3–16 (the San Carlos Agent) ☆. Virology 36:201–211.

10. Fender P, Ruigrok RW, Gout E, Buffet S, Chroboczek J. 1997. Adenovirus dodecahedron, a new vector for human gene transfer. Nature Biotechnology 15:52–56.

11. Liew MWO, Chuan YP, Middelberg APJ. 2012. High-yield and scalable cell-free assembly of virus-like particles by dilution. Biochemical Engineering Journal 67:88–96.

12. Boulanger PA, Puvion F. 1976. Occurrence of a peculiar type of adenovirus 2 penton oligomer. Intervirology 7:126–134.

13. Wilcox WC, Ginsberg HS. 1963. STRUCTURE OF TYPE 5 ADENOVIRUS. 118:307–314.

14. Karayan L, Gay B, Gerfaux J, Boulanger PA. 1994. Oligomerization of recombinant penton base of adenovirus type 2 and its assembly with fiber in baculovirus-infected cells. Virology 202:782–795.

15. Fender P, Schoehn G, Foucaudgamen J, Gout E, Garcel A, Drouet E, Chroboczek J. 2003. Adenovirus Dodecahedron Allows Large Multimeric Protein Transduction in Human Cells. Journal of Virology 77:4960–4964.

16. Garcel A, Gout E, Timmins J, Chroboczek J, Fender P. 2006. Protein transduction into human cells by adenovirus dodecahedron using WW domains as universal adaptors. Journal of Gene Medicine 8:524–531.

17. Zochowska M, Paca A, Schoehn G, Andrieu JP, Chroboczek J, Dublet B, Szolajska E. 2009. Adenovirus dodecahedron, as a drug delivery vector. Plos One 4:e5569–e5569.

18. Naskalska A, Szolajska E, Chaperot L, Angel J, Plumas J, Chroboczek J. 2009. Influenza recombinant vaccine: matrix protein M1 on the platform of the adenovirus dodecahedron. Vaccine 27:7385–7393.

19. Naskalska A, Szolajska E, Chaperot L, Angel J, Plumas J, Chroboczek J. 2009. Influenza recombinant vaccine: Matrix protein M1 on the platform of the adenovirus dodecahedron. Vaccine 27:7385–7393.

20. Villegas-Mendez A, Garin MI, Pineda-Molina E, Veratti E, Bueren JA, Fender P, Lenormand J-L. 2010. In Vivo Delivery of Antigens by Adenovirus Dodecahedron Induces Cellular and Humoral Immune Responses to Elicit Antitumor Immunity. Molecular Therapy 18:1046–1053.

21. Kang X, Xiao H-H, Song H-Q, Jing X-B, Yan L-S, Qi R-G. 2015. Advances in drug delivery system for platinum agents based combination therapy. Cancer Biology & Medicine 12:362–374.

22. Mccracken RM, Mcferran JB, Evans RT, Connor TJ. 1976. Experimental studies on the aetiology of inclusion body hepatitis. Avian Pathology Journal of the Wvpa 5:325–339.

23. Hess M. 2000. Detection and differentiation of avian adenoviruses: A review. Avian Pathology 29:195–206.

24. Wang X, Tang Q, Chu Z, Wang P, Luo C, Zhang Y, Fang X, Qiu L, Dang R, Yang Z. 2017. Immune protection efficacy of FAdV-4 surface proteins fiber-1, fiber-2, hexon and penton base. Virus Research 245:1.

25. Schachner A, Marek A, Jaskulska B, Bilic I, Hess M. 2014. Recombinant FAdV-4 fiber-2 protein protects chickens against hepatitis-hydropericardium syndrome (HHS). Vaccine 32:1086–1092.

26. You GJ, Xia J, Yao KC, Liu P, Zhao Q, Cao SJ, Li SY, Huang XB, Han XF, Wu YPWR. 2017. Isolation and molecular characterization of prevalent Fowl adenovirus strains in southwestern China during 2015–2016 for the development of a control strategy. Emerging Microbes & Infections 6:e103.

27. Changjing L, Haiying L, Dongdong W, Jingjing W, Youming W, Shouchun W, Jida L, Ping L, Jianlin W, Shouzhen X. 2016. Characterization of fowl adenoviruses isolated between 2007 and 2014 in China. Veterinary Microbiology 197:62–67.

28. Zhang T, Jin Q, Ding P, Wang Y, Chai Y, Li Y, Liu X, Luo J, Zhang G. 2016. Molecular epidemiology of hydropericardium syndrome outbreak-associated serotype 4 fowl adenovirus isolates in central China. Virology Journal 13:188.

29. Kim MS, Lim TH, Lee DH, Youn HN, Yuk SS, Kim BY, Choi SW, Jung CH, Han JH, Song CS. 2014. An inactivated oil-emulsion fowl Adenovirus serotype 4 vaccine provides broad cross-protection against various serotypes of fowl Adenovirus. Vaccine 32:3564–3568.

30. Kumar R, Chandra R, Shukla SK, Agrawal DK, Kumar M. 1997. Hydropericardium syndrome (HPS) in India: A preliminary study on the causative agent and control of the disease by inactivated autogenous vaccine. Tropical Animal Health & Production 29:158.

31. Anjum AD. 1990. Experimental transmission of hydropericardium syndrome and protection against it in commercial broiler chickens. Avian Pathology 19:655–660.

32. Schonewille E, Jaspers R, Paul G, Hess M. 2010. Specific-pathogen-free chickens vaccinated with a live FAdV-4 vaccine are fully protected against a severe challenge even in the absence of neutralizing antibodies. Avian Diseases 54:905–910.

33. Mansoor MK, Hussain I, Arshad M, Muhammad G. 2011. Preparation and evaluation of chicken embryo-adapted fowl adenovirus serotype 4 vaccine in broiler chickens. Trop Anim Health Prod 43:331–338.

34. Gupta A, Ahmed KA, Ayalew LE, Popowich S, Kurukulasuriya S, Goonewardene K, Gunawardana T, Karunarathna R, Ojkic D, Tikoo SK. 2017. Immunogenicity and protective efficacy of virus-like particles and recombinant fiber proteins in broiler-breeder vaccination against fowl adenovirus (FAdV)-8b. Vaccine 35:2716.

35. Wang X, Tang Q, Chu Z, Wang P, Luo C, Zhang Y, Fang X, Qiu L, Dang R, Yang Z. 2018. Immune protection efficacy of FAdV-4 surface proteins fiber-1, fiber-2, hexon and penton base. Virus Research 245:1–6.

36. Wang X, Wang X, Jia Y, Wang C, Tang Q, Han Q, Xiao S, Yang Z. 2017. Coadministration of Recombinant Adenovirus Expressing GM-CSF with Inactivated H5N1 Avian Influenza Vaccine Increased the Immune Responses and Protective Efficacy Against a Wild Bird Source of H5N1 Challenge. Journal of interferon & cytokine research: the official journal of the International Society for Interferon and Cytokine Research 37:467–473.

37. Wold WSM, Toth K. 2013. Adenovirus Vectors for Gene Therapy, Vaccination and Cancer Gene Therapy. Current gene therapy 13:421–433.

38. Castello R, Borzone R, D’Aria S, Annunziata P, Piccolo P, Brunetti-Pierri N. 2015. Helper-dependent adenoviral vectors for liver-directed gene therapy of primary hyperoxaluria type 1. Gene Therapy 23:129.

39. Norrby E. 1968. Comparison of Soluble Components of Adenovirus Types 3 and 11. Journal of General Virology 2:135–142.

40. Fender P, Boussaid A, Mezin P, Chroboczek J. 2005. Synthesis, cellular localization, and quantification of penton-dodecahedron in serotype 3 adenovirus-infected cells. Virology 340:167–173.

41. Vivès RR, Lortat-Jacob H, Chroboczek J, Fender P. 2004. Heparan sulfate proteoglycan mediates the selective attachment and internalization of serotype 3 human adenovirus dodecahedron. Virology 321:332–340.

42. Fender P, Schoehn G, Perron-Sierra F, Tucker GC, Lortat-Jacob H. 2008. Adenovirus dodecahedron cell attachment and entry are mediated by heparan sulfate and integrins and vary along the cell cycle. Virology 371:155–164.

43. Li T-C, Takeda N, Kato K, Nilsson J, Xing L, Haag L, Cheng RH, Miyamura T. 2003. Characterization of self-assembled virus-like particles of human polyomavirus BK generated by recombinant baculoviruses. Virology 311:115–124.

44. Song L, Nakaar V, Kavita U, Price A, Huleatt J, Tang J, Jacobs A, Liu G, Huang Y, Desai P, Maksymiuk G, Takahashi V, Umlauf S, Reiserova L, Bell R, Li H, Zhang Y, McDonald WF, Powell TJ, Tussey L. 2008. Efficacious Recombinant Influenza Vaccines Produced by High Yield Bacterial Expression: A Solution to Global Pandemic and Seasonal Needs. PLoS ONE 3:e2257.

45. Shi L, Sanyal G, Ni A, Luo Z, Doshna S, Wang B, Graham TL, Wang N, Volkin DB. 2005. Stabilization of human papillomavirus virus-like particles by non-ionic surfactants. Journal of Pharmaceutical Sciences 94:1538–1551.

46. Zochowska M, Paca A, Schoehn G, Andrieu J-P, Chroboczek J, Dublet B, Szolajska E. 2009. Adenovirus Dodecahedron, as a Drug Delivery Vector. PLOS ONE 4:e5569.

47. Gupta A, Ahmed KA, Ayalew LE, Popowich S, Kurukulasuriya S, Goonewardene K, Gunawardana T, Karunarathna R, Ojkic D, Tikoo SK. 2017. Immunogenicity and protective efficacy of virus-like particles and recombinant fiber proteins in broiler-breeder vaccination against fowl adenovirus (FAdV)-8b. Vaccine 35:2716–2722.

48. Schachner A, Marek A, Jaskulska B, Bilic I, Hess M. 2014. Recombinant FAdV-4 fiber-2 protein protects chickens against hepatitis–hydropericardium syndrome (HHS). Vaccine 32:1086–1092.

49. Wang X, Qiu L, Hao H, Zhang W, Fu X, Zhang H, He S, Zhang S, Du E, Yang Z. 2012. Adenovirus-based oral vaccine for rabbit hemorrhagic disease. Veterinary Immunology & Immunopathology 145:277–282.

